# DIMPLE: AN R PACKAGE TO QUANTIFY, VISUALIZE, AND MODEL SPATIAL CELLULAR INTERACTIONS FROM MULTIPLEX IMAGING WITH DISTANCE MATRICES

**DOI:** 10.1101/2023.07.20.548170

**Authors:** Maria Masotti, Nathaniel Osher, Joel Eliason, Arvind Rao, Veerabhadran Baladandayuthapani

**Affiliations:** UNIVERSITY OF MICHIGAN DEPARTMENT BIOSTATISTICS; UNIVERSITY OF MICHIGAN DEPARTMENT OF COMPUTATION MEDICINE AND BIOINFORMATICS

**Author notes:** THESE AUTHORS CONTRIBUTED EQUALLY (Maria Masotti, Nathaniel Osher, Joel Eliason).

**Keywords:** Multiplex Imaging, Point Process, Spatial Statistics

## Abstract

The tumor microenvironment (TME) is a complex ecosystem containing tumor cells, other surrounding cells, blood vessels, and extracellular matrix. Recent advances in multiplexed imaging technologies allow researchers to map several cellular markers in the TME at the single cell level while preserving their spatial locations. Evidence is mounting that cellular interactions in the TME can promote or inhibit tumor development and contribute to drug resistance. Current statistical approaches to quantify cell-cell interactions do not readily scale to the outputs of new imaging technologies which can distinguish many unique cell phenotypes in one image. We propose a scalable analytical framework and accompanying R package, DIMPLE, to quantify, visualize, and model cell-cell interactions in the TME. In application of DIMPLE to publicly available MI data, we uncover statistically significant associations between image-level measures of cell-cell interactions and patient-level covariates.

## 1. The bigger picture

The `DIMPLE` R package fills a major gap for conducting spatial analysis of multiplex tissue imaging (MI) data. Existing approaches to quantifying spatial cell-cell interactions in the tumor microenvironment (TME) are based on spatial metrics that are computationally expensive and present inferential and interpretational challenges. `DIMPLE` was designed specifically for MI single cell data. It is robust to holes and heterogeneity of spatial distributions of the cells. It can compute all pairwise cellular interactions in parallel and display them in an easy-to-interpret matrix. These matrices can be passed as an outcome or covariate in modeling for testing association with patient-level clinical metadata. `DIMPLE` provides an end-to-end pipeline to quantify, visualize and model spatial cellular interactions which will enable discovery of novel associations of inter-cellular interactions in the TME with patient outcomes.

## 2. Introduction

The tumor microenvironment (TME) refers to the complex network of cells and other structures surrounding a tumor. This microenvironment is essential for the growth, survival, and metastasis of cancer cells.^1^ Understanding the mechanisms underlying the TME is crucial for developing new and more effective cancer therapies.^2, 3^ Recent breakthroughs in multiplex tissue imaging (MI) allow researchers to simultaneously visualize and quantify multiple biomarkers in a single tissue sample while preserving their spatial information.^4^ Specific MI technologies include PhenoImager, formerly known as Vectra, PhenoCycler, formerly known as CO-Detection by Indexing (CODEX), Multiplexed ion beam imaging by time-of-flight (MIBI), Imaging mass cytometry (IMC), Matrix-assisted laser desorption ionization mass spectrometry imaging (MALDI-MSI), and Digital spatial analysis (GeoMx/DSP/CosMx). These technologies produce high resolution maps of multiple functional and phenotypic markers on a single tissue section. Each single cell in the tissue can be phenotyped based on the marker intensities. One of the key analytical goals for high-resolution images from these technologies is to understand the interactions between different cells in the TME and how they contribute to tumor growth, metastasis, and drug resistance.

Spatial analysis of MI-derived data may offer insight into how cellular crosstalk and heterogeneity affect cancer prognoses and responses to treatment. Accurately characterizing such interactions and heterogeneity in the TME – both within and across tumor samples – is essential in deciphering cellular mechanisms that underlie cancer evolution and development. Several recent studies discovered novel cellular interactions in the TME. A MI study on the lung adenocarcinoma TME in 153 patients with resected tumors found that expression of major histocompatibility complex II (MHCII) associates with tumor and immune interaction within the TME.^5^ This suggests that cancer cell-specific expression of MHCII may represent a biomarker for the immune system’s recognition and activation against the tumor. An MI study of ovarian cancer found that the proximity between tumor-associated macrophages and B cells or CD4 T cells significantly correlated with overall survival.^6^ An MI study of tissue samples from the colorectal cancer (CRC) invasive front found that colocalization of PD-1 postive CD-4 positive T cells with granulocytes was positively correlated with survival in a high-risk patient subset.^7^

Despite structurally similar data, each of these studies attempts to quantify cellular interaction with profoundly different techniques. In the study of the lung cancer TME,^5^ Euclidean distances between individual tumor cells and the nearest immune cell phenotypes were calculated. In the study of ovarian cancer,^6^ an interaction variable based on the number of one cell type within a certain radius of another was calculated. This measure was binned and used downstream in survival outcome modeling. In the study of CRC,^7^ a complex clustering procedure was used to partition each image into a neighborhood. The neighborhoods were roughly defined by prevalence of certain cells. Then summary statistics were computed in each neighborhood and compared between neighborhoods and images. These studies demonstrate a lack of consensus on an approach for measuring cellular interaction in the TME. Measures of interaction based at the cell-level, such as Ripley’s K,^8^ Besag’s L,^9^ Marcon’s M^10^ and nearest neighbor distance^11^ may not readily scale with increasing number of cell types or number of images. Specifically, pairwise interactions grow quadratically with the number of cell types, so discovery driven approaches can quickly become computationally infeasible. These cell-level measures are functions of a radius (*r*), which must be pre-specified or tuned in order to calculate a univariate measure of interaction for downstream analysis. Alternatively, more complex modeling procedures are required to study the heterogeneity of cell-cell interactions using these functions over a range of *r* as functional data inputs.^12^ The values of these measures are not interpretable alone and require comparison with the function estimated under complete spatial randomness (CSR). The value of the function under CSR is highly sensitive to spatial inhomogeneity, uneven distribution of cells across the image, and “holes” or areas of missing data.^13^ Unfortunately, these features are common in MI data, so calculation of the CSR must be performed by permutation for each individual image, which can, again, be quite computationally expensive. Software tools have been developed to compute cell-level measures of interaction. One of the most widely used, the R package “spatstat”,^14^ is a comprehensive set of functions for spatial point patterns. The R packages “spatialTIME”^15^ and “spicyR”^16^ were designed specifically to store and analyze data from multiplex imaging. These software packages compute some of the cell-level measures mentioned above and utilize permutation-based approaches to compare them with CSR. For the purpose of driving discovery from this large and complex data, an interpretable measure of cell-cell interaction and software that scales to current and future MI outputs is needed.

To address these challenges, we present `DIMPLE`: Distance Matrices for Multiplex Imaging, along with an R package and accompanying R Shiny app, filling the gap for a scalable and statistically-savvy data science toolkit for researchers conducting MI experiments (see Figure 1). Briefly, `DIMPLE` first computes a spatially smooth non-parametric kernel density estimate (KDE) of each cell type for each image, the parameters of which are user-defined. Then, a distance or similarity metric is applied to each unique pairwise combination of the KDEs for each image. These univariate distance metrics are organized in matrix form and can be visualized using heatmaps or networks. Users have the option to attach patient-level metadata, allowing users to test associations between aggregated distance metrics and patient covariates or outcomes. `DIMPLE` is designed to accommodate data from a variety of sources, requiring only image IDs, spatial coordinates, and a cell type or marker status for each cell. The plotting functions and accompanying Shiny app generate publication-quality figures to visualize heterogeneity in the cell-cell interactions across patient-level covariates. Our framework is computationally efficient and requires no permutations to interpret measures of cellular interaction. The image-level and optional patient-level data along with computed intensities and distance matrices are organized with all relevant information in an R S3 object. `DIMPLE` is designed to appeal to novice R users, but outputs from the package can be used downstream in more complex statistical analysis such as survival analysis, mixed effects modeling, clustering, and variable selection.

**Figure 1.**
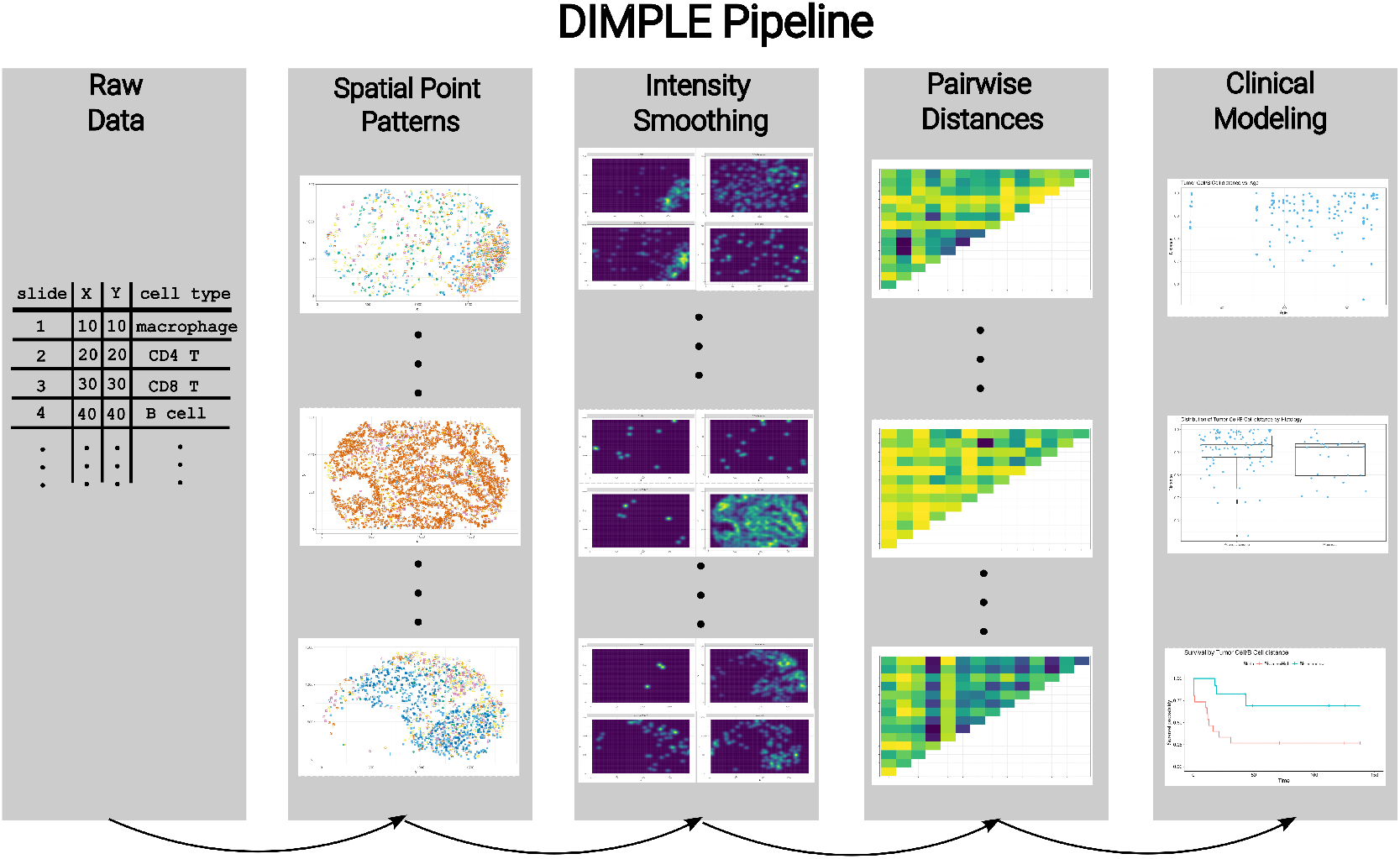
Overview of `DIMPLE` pipeline, from raw data to clinical analysis. First, data is supplied to `DIMPLE` in a simple format. Second, a point pattern representation of the data is generated for each image. Third, cell type intensity surfaces are estimated. Fourth, distances are computed from the intensities for each pair of cell types. Finally, pairwise distances may be easily used downstream in statistical analyses in combination with patient metadata.

To illustrate the functionality of the `DIMPLE` software, we will utilize publicly available MI data of the lung adenocarcinoma TME in 153 patients with resected tumors from Johnson et al.^5^ In Section 3, we describe each of the analysis steps (columns of Figure 1) using this data as an example. Results from our application of `DIMPLE` to this data reveals statistically significant associations between different types of immune cells and CK+ tumor cells. These findings agree with findings from Johnson et al,^5^ who found that expression of major histocompatibility complex II (MHCII) associates with tumor and immune interaction within the TME. We conclude Section 3 with a tutorial for generating `DIMPLE` distance matrices at quantile-specific partitions of the MI data in Section 3.5, an overview of our R Shiny App in Section 3.6, and a note on simulating MI data in Section 3.7.

## 3. Results

In this section, we describe how to use the `DIMPLE` package and provide complete coding examples. The data can be downloaded using the `VectraPolarisData` package.^17^ See the supplemental materials for the R code used to download and process the data for use in `DIMPLE`. The overall analyses pipeline is organized into 4 main steps. Step 1 entails reading in data and storing it as a `MltplxExperiment`. Step 2 entails generating intensity surface estimates for each cell type and image in the dataset. Step 3 entails generating distance matrix representations of cellular interaction for each image. Step 4 entails using the pairwise distances in downstream analysis. This section concludes with a tutorial of generating quantile-specific distance matrices to investigate patterns of cellular interaction at various partitions of the image.

**Figure 2.**
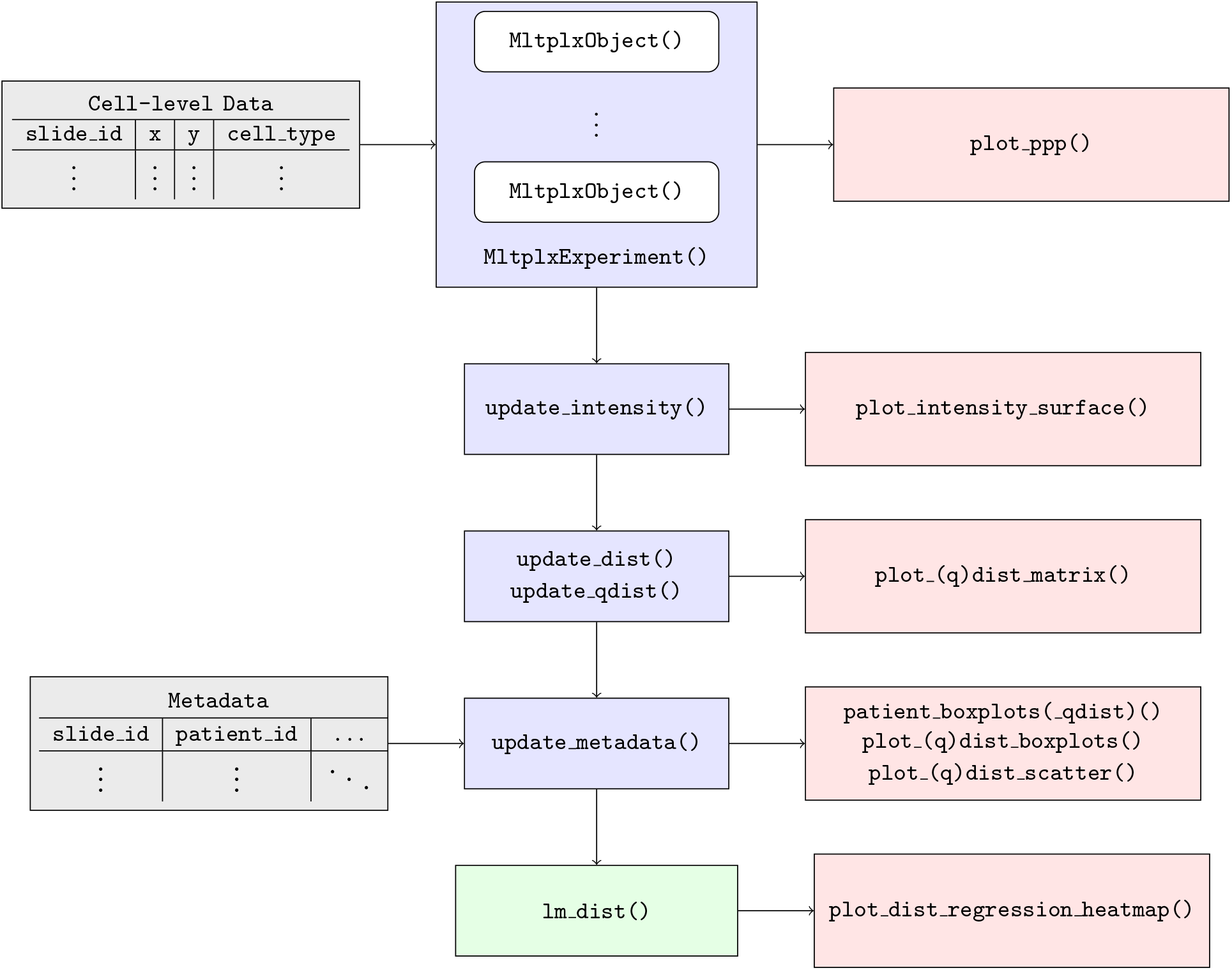
A flowchart representing inputs (grey), infrastructure functions (blue), visualization functions (red), and inference functions (green).

### 3.1. Step 1: Input data and construct

`MltplxExperiment`. To generate measures of cellular interaction using the `DIMPLE` package, the minimal amount of information needed is four vectors with length equal to the total number of cells in the multiplex imaging dataset:

1. x, the x-coordinates of each cell
2. y, the y-coordinates of each cell
3. `marks`, the cell-type of each cell
4. `slide id`, the ID of the image that each cell is from

The `new MltplxExperiment`function takes at least those four inputs: `x`, `y`, `marks`, and `slide_id`. It returns an S3 object of class `MltplxExperiment`. This object is at its core a list of all of the slides in the data set, each with its corresponding cell types and locations stored for easy access, along with additional metadata. In fact, it can be indexed the way a standard list is indexed, using single- and double-bracket expressions as demonstrated below.

We will store the example data in a `MltplxExperiment` called `lung_experiment`:

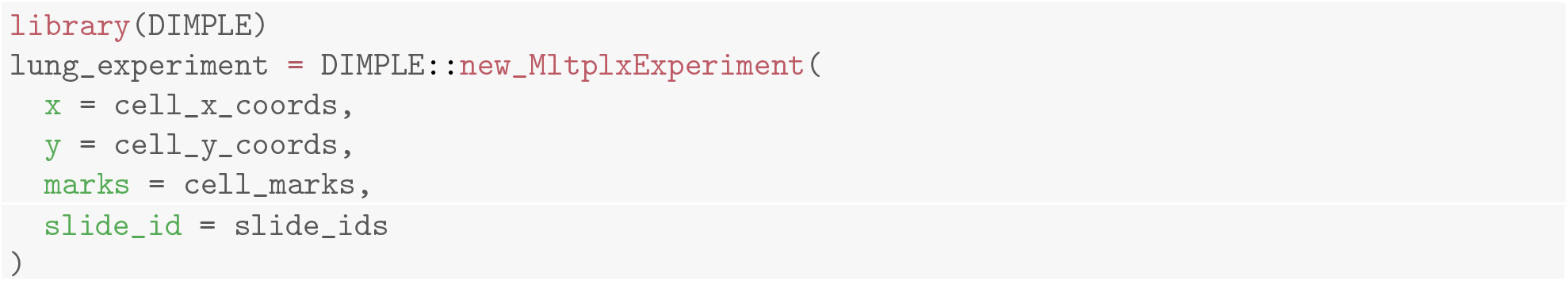

Each slide in the dataset is stored as a `MltplxObject` within the `MltplxExperiment` object. A `MltplxObject` represents a labeled collection of cell types and locations for each slide in the data set. The S3 method `plot` is implemented for `MltplxObject`, which allows for quick inspection for a given slide:

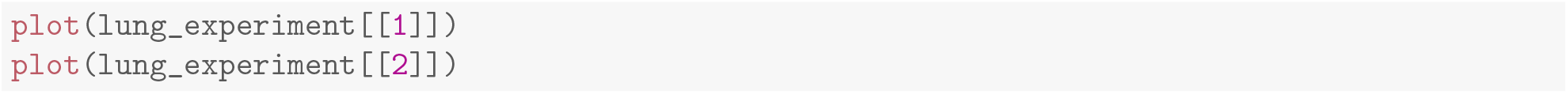

**Figure 3.**
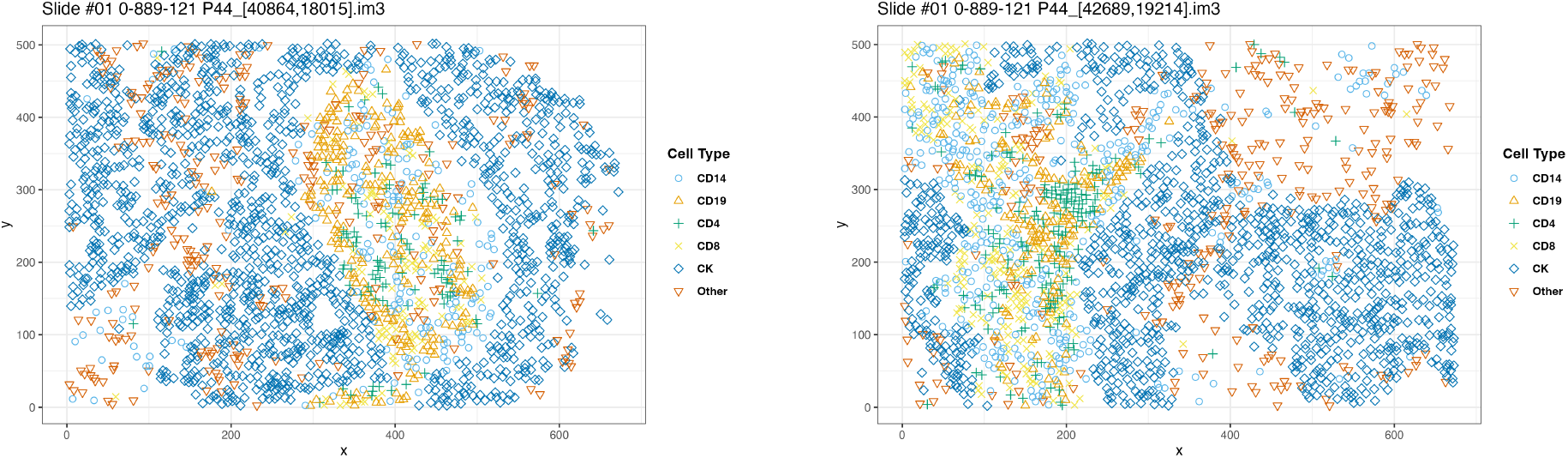
A point pattern plot of the first (left) and second (right) image contained in `lung_experiment` generated with the `plot()` function. Combinations of color and shape represents the cell phenotype. These images come from the same patient, revealing heterogeneity within patient and over space - both hallmarks of MI data.

### 3.2. Step 2: Generating and visualizing cell type intensities

The intensity of a point process can be thought of as an average, or first-moment. The intensity function can be estimated non-parametrically by kernel estimation. The `DIMPLE` package uses an isotropic Gaussian kernel and Diggle’s correction^18, 19^ to reduce the bias from edge effects. There are two arguments necessary to generate cell intensities: `ps` and `bw`. `ps` controls the size of the grid at which the intensity estimates will be generated. `ps` is short for “pixel size,” since the resulting intensities can often resemble highly pixelized images. `bw` controls the smoothing of the resulting pixels, i.e. the degree to which the values of certain pixels should resemble their neighbors. Larger values of `bw` will result in “smoother” intensity surfaces. The smoothing bandwidth can be tuned by the user and should be chosen with some prior biological knowledge about the “radius of influence” that a given cell type has within a tissue. `DIMPLE` makes it trivial to generate intensity grids for points of all types for entire `MltplxExperiment` objects as well as individual `MltplxObject` objects. To expedite computations, users may implement intensity smoothing and calculation of distances in parallel by uncommenting the second line of code below:

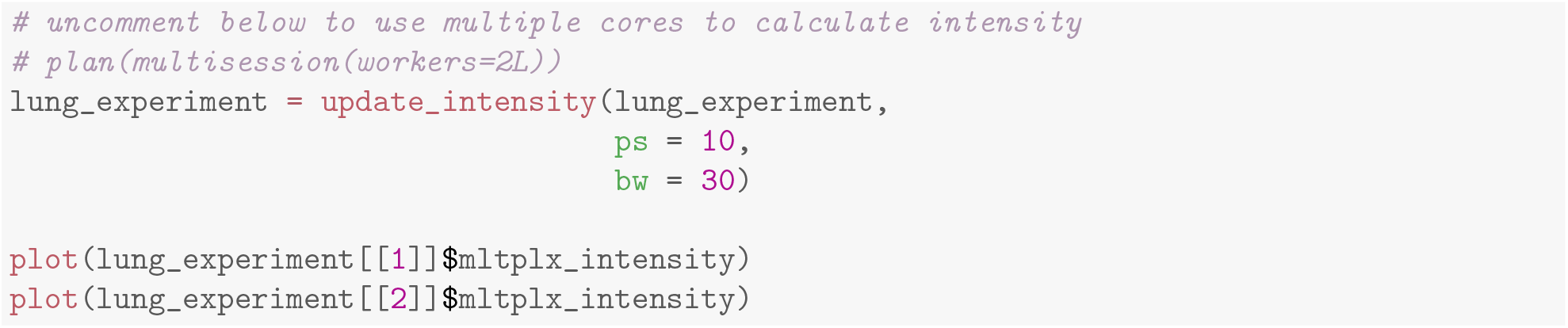

### 3.3. Step 3: Generating and visualizing pairwise distance matrices

Having generated appropriate intensity functions, a reasonable question to ask is: how *similar* or *different* are the distributions of cells of different types within a given slide? For example, the intensity surfaces in the previous example seem to suggest that, for this specific patient, while there is considerable overlap between the positions of CD4 positive and CD19 positive cells, there is very little overlap between either of these types and CK positive cells.

**Figure 4.**
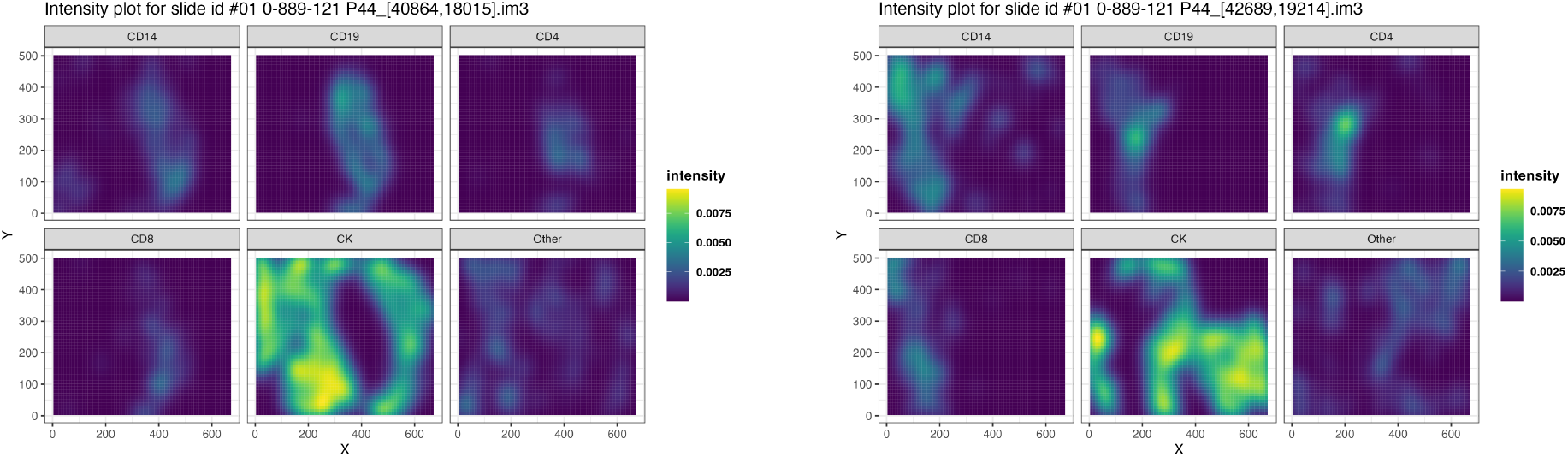
Estimated intensity surfaces for each cell phenotype from the two point patterns in Figure 3 generated using the code above. Color represents intensity with brighter color indicating greater intensity values.

To formalize this intuition in a way that can allow comparison across different slides, one can employ measures of distance between the intensities of different cell types within a given slide. Although, any user defined distance metric can be used, we prefer Jensen-Shannon distance (JSD) which is a method of measuring similarity between two probability distributions, as discussed in.^20^ It is a symmetrized and smoothed version of the Kullback-Leibler divergence (KLD). The Jensen-Shannon distance (JSD) takes the square-root of the Jensen-Shannon divergence so that it fulfills the axioms of a metric. JSD is our preferred metric for this task because it is bounded by 0 (perfect overlap) and 1 (complete separation), symmetric, and does not suffer from inflation due to the presence of zeros (holes in the image). The JSD is defined by:

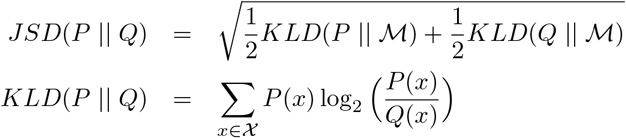

where *P* and *Q* are probability distributions defined on the same sample space *χ* and 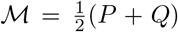. Having computed intensities for each cell type in a `MltplxExperiment` object, pairwise distances can be added easily by specifying a distance metric. Here we use `jsd`:

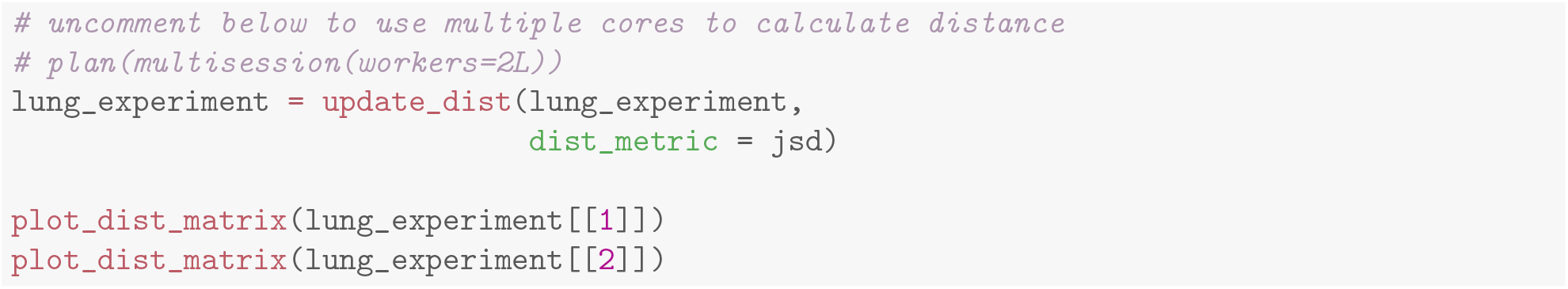

The `dist_metric` can be user-defined. Any function that takes in two vectors of the same length as an argument and produces a single scalar can be used in place of `jsd`.

The resulting distance matrices formalize our previous intuition. Since higher values of Jensen-Shannon distance indicate more different distributions (and vice versa for smaller values), it does indeed seem that there is quite a bit of overlap between the CD19 positive and CD4 positive cells, while the distributions of these cells are quite far from those of the CK positive cells in the two images within this patient. Boxplots of image-level Tumor to CD8 distances reveal significant within- and between-patient heterogeneity, see Figure 6.

**Figure 5.**
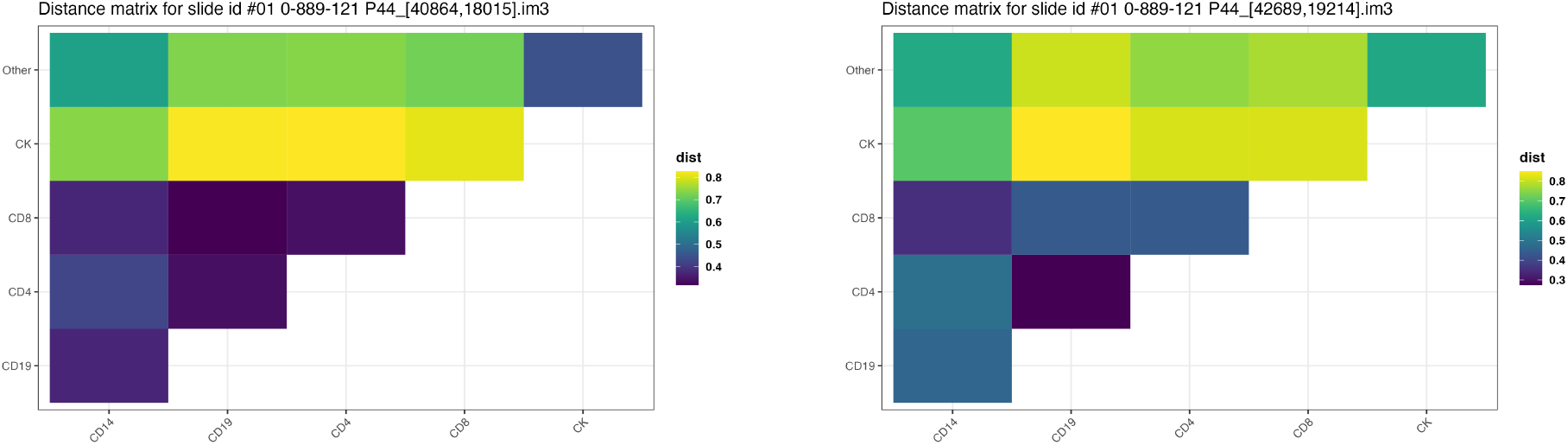
Estimated pairwise distances for the point patterns in Figure 3 generated using the code above. Color represents JSD with brighter color indicating greater distance.

**Figure 6.**
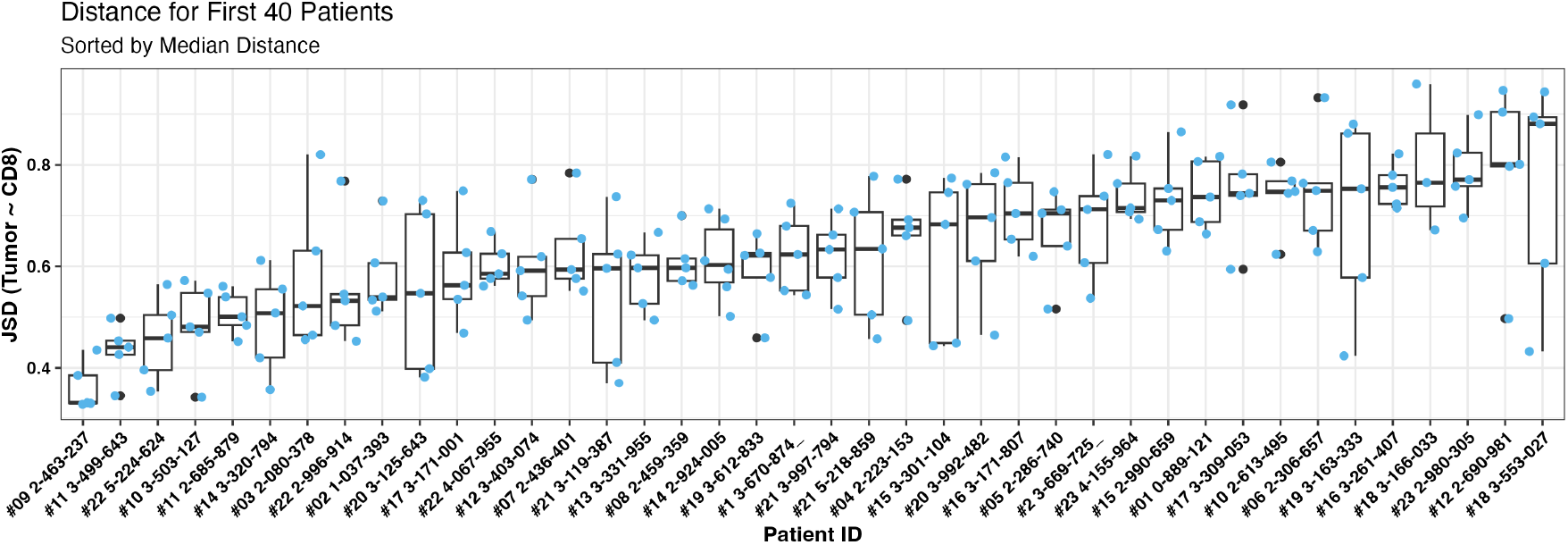
Distribution of biopsy distances between Tumor cells and CD8 T cells within and between of first forty patients in the data set, sorted by median distance.

We conclude this section by noting that the changes made to the `lung_experiment` object can be discerned at a glance by simply invoking the `print` function on the object, implicitly or explicitly:

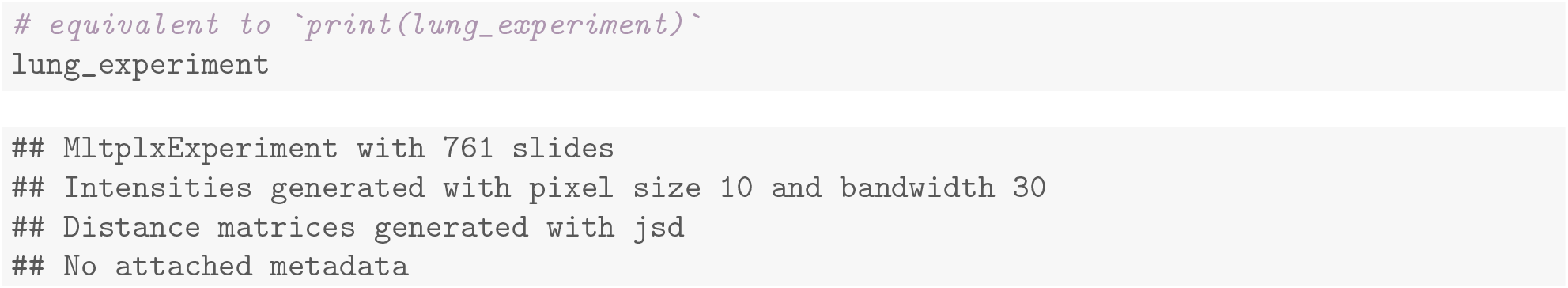

### 3.4. Step 4: Adding Patient Metadata, Visualizing, and Modeling Associations

Patient level data can also be stored in the `MltplxExperiment` for the purpose of visualizing and modeling cellular interactions with patient outcomes. Patient metadata can be attached to the `lung_experiment` object using the `update_metadata` function. The patient metadata must contain 2 columns and each row corresponds to a `slide_id` in the `MltplxExperiment`:

1. `slide id` contains the same set of IDs as the `MltplxExperiment`
2. `patient id` links the `slide_id` to the patient identifier

The metadata can be added to `lung_experiment` via:

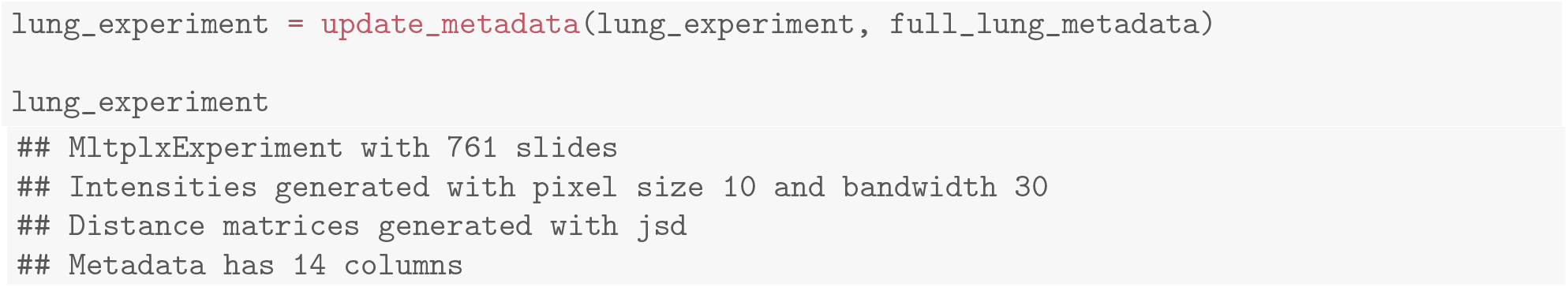

The generic function `as tibble` facilitates downstream modeling and analysis by generating a tibble containing all of the columns that were already present in the metadata and a column for each pair of cell types. Users may easily include the pairwise distances as covariates in survival or regression modeling of patient outcomes, model them as outcomes in a mixed effects model, or uncover natural groupings of the images through unsupervised clustering.

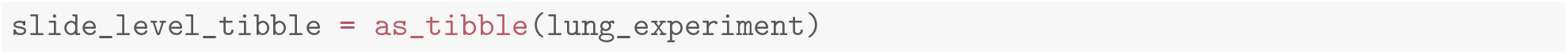

Basic linear regression can be performed using the `lm dist` function which fits separate models for each pairwise distance to test for association with a variable from the patient metadata. In this example, “mchII status” is a column of the patient metadata and indicates whether a patient’s tumor cells expressed a high or low amount of major histocompatibility complex II. The user can specify variables to adjust for and aggregating functions to use. The default settings adjust for cell type counts and aggregates images within patient via the median. The tested coefficient for each model can be displayed graphically using `plot_dist_regression heatmap`.

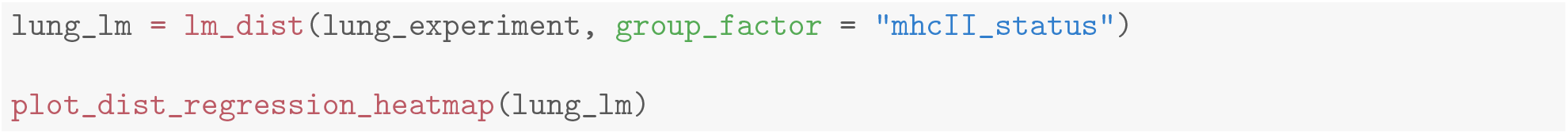

**Figure 7.**
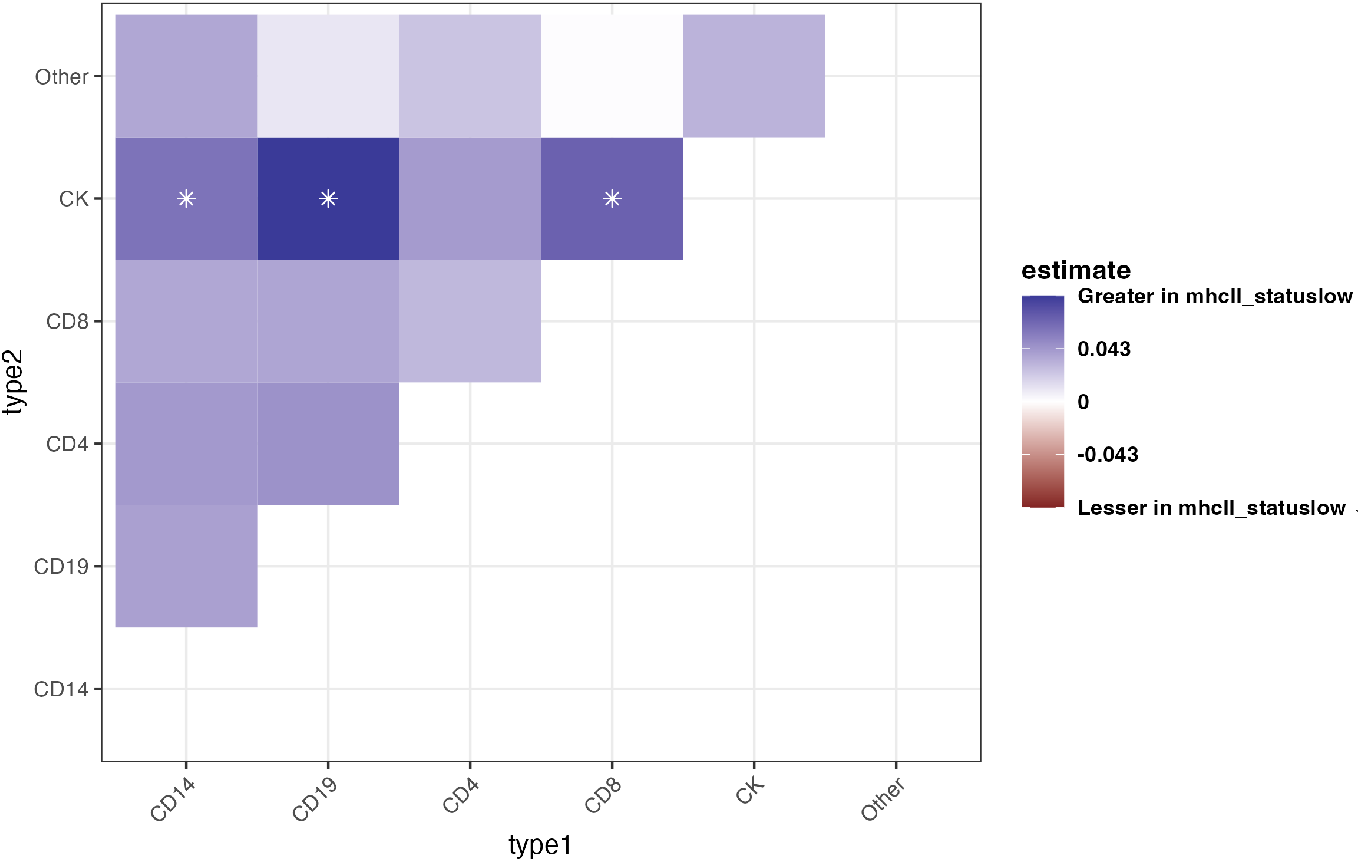
Estimated effect of mhcII status low on pairwise cellular distances. Blue indicates a positive effect on distance and red indicates a negative effect on distance. Stars indicate fdr adjusted p-values less than 0.05.

**Figure 8.**
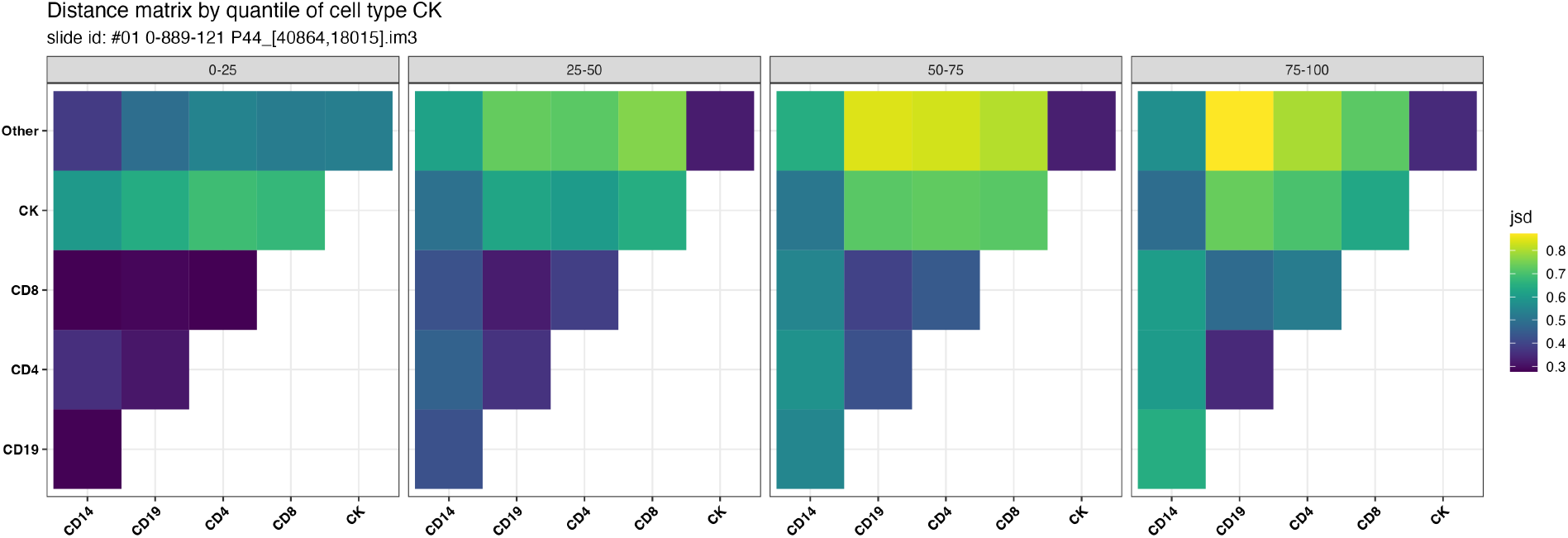
Quantile specific distance matrices generated by the code above for the point pattern in figure 3 faceted by intensity of CK+ cells. The color represents JSD with brighter color indicating greater distance.

**Figure 9.**
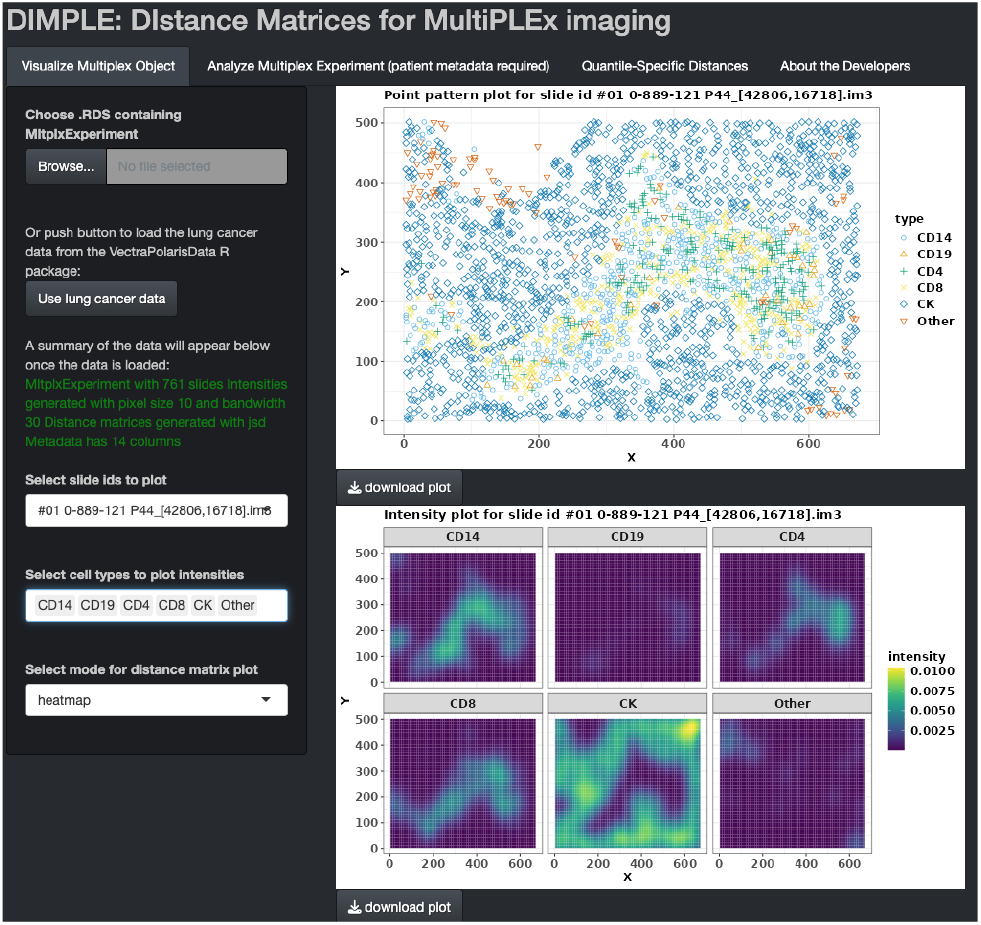
The R Shiny app allows users to explore and visualize MI data. Users can simply upload a `MltplxExperiment` object or explore the lung data referenced in this article with the click of a button.

In this example data, low expression of MCHII is associated with decreased infiltration of immune cells with CK positive tumor cells. This finding is in agreement with Johnson et al.^5^

As an alternative to aggregating distances within patient, distances may be modeled as outcomes in mixed-effect models with patient specific random effects to account for within-patient correlations.

### 3.5. Generating and visualizing quantile-specific distance matrices

Patient outcomes or covariates may not be associated with pairwise cellular interactions on the scale of the entire image. Rather, cellular interaction at different regions of the image may be associated with outcomes. The distance matrices may be computed at various partitions of the image. We have implemented functionality to compute distance matrices in regions defined by user-defined quantiles of one cell-type intensity. The `update_qdist` function takes the following arguments:

1. `dist metric` determines the distance metric
2. `mask type` determines by which cell type intensity to partition
3. `q probs` data frame represents a range of quantiles to partition the intensity by

The following code partitions each image in `lung_experiment` into distinct areas based on quartiles of the intensity of CK positive cells:

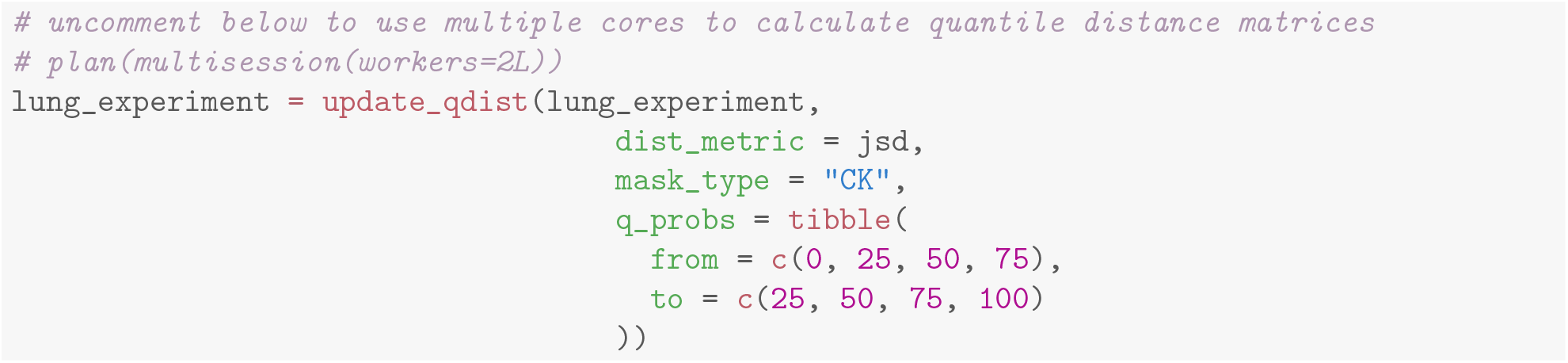

The quantile-specific distance matrices can be visualized using the `plot qdist` function:

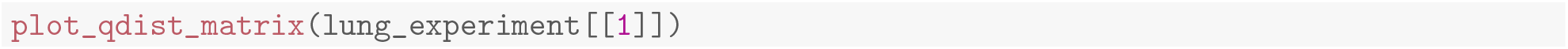

Across the different quartiles, the patterns of spatial interactions between different cell types can vary considerably. In this particular image, as the intensity of “CK” phenotyped cells increases, the distance between the distributions of many of the various immune phenotyped cells also increases. Interestingly, though, the distance between CD14 phenotyped cells and CK phenotyped cells decreases. Varying levels of immune cell co-localization across regions of the tumor may indicate increasing concentrations of cytokines or other signaling molecules in the core tumor regions.^21^ Quantile-specific distance matrices generated by `DIMPLE` allow one to probe and explore the natural heterogeneity that arises from such multifactorial and complex interactions in the TME.

### 3.6. Using the DIMPLE Shiny App

The R Shiny app is an additional tool to help researchers generate publication ready plots using the outputted distance matrices and intensities from the R package. Plots included in this proposal may be generated with this app. It produces color-blind friendly visualizations of individual intensities and distance matrices, aggregated data, and model outputs. Each plot may be downloaded as a pdf file. The app is freely available at: https://bayesrx.shinyapps.io/DIMPLE_Shiny/. The Shiny app accepts a `MltplxExperiment` saved as a `.RDS` data file. Or, a user can explore the lung cancer data referenced in this tutorial with the click of a button. The Shiny app is organized into the following 3 pages:

1. **Visualize Multiplex Object** visually explore individual `MltplxObjects` by selecting a slide ID.
2. **Analyze Multiplex Experiment** basic inference and visualization of model outputs in combination with patient-level metadata
3. **Quantile-Specific Distances** explore, visualize, and make inference on quantile-specific distance matrices

### 3.7. Simulation of MI data

Finally, we have developed several functions to simulate MI data from intensity surfaces. These functions could be used in testing how different parameters of the intensity smoothing effect measures of cell-cell interactions. We provide a tutorial of those functions in Section 2 of the Supplemental Materials.

## 4. Discussion

We introduce `DIMPLE`, a statistical software package designed to probe the relationships between cell types obtained from MI experiments. `DIMPLE` converts spatial and phenotypic information captured in MI assays to continuously-varying non-parametric kernel density estimates (KDEs) of the point process intensity function with a user-defined smoothing bandwidth. Then, a user-defined distance metric is applied to the pairwise combinations of the KDEs for each cell type.

The resulting distance matrix can be further explored and, when combined with patient-level metadata, can be used to identify potential biomarkers wthin the TME. `DIMPLE` was developed primarily to assist in exploratory work, especially in developing new hypotheses around second-order cell type relationships. It contains many diverse plotting functions, each of which is useful for developing intuition on different aspects of first- and second-order cell type relationships in isolation or with respect to covariates. In addition, `DIMPLE` includes simple but powerful simulation functionality that allows the user to test how different parameters will change downstream outcomes; for details see Section 2 of the supplementary materials.

Perhaps most importantly, all of this functionality is built around a single simple, flexible, and easily extendable data structure: the `MltplxExperiment` object. One of the more banal challenges of working with MI data is the many different forms the data may take during the analysis. One may need to convert and translate between lists of `ppp` objects, data frames of patient data, and matrices of different metrics, all to analyze a single data set. This does not even begin to explore the challenges of adapting such code to data from different platforms and with different markers. This is, to some degree, unavoidable-`DIMPLE` certainly utilizes a number of different data structures “under the hood” to store data. But by creating an extendable and platform-agnostic class that both standardizes the form in which data is stored and takes care of conversions and translations as necessary, `DIMPLE`minimizes the need to write “boilerplate” code and maximizes the ability to write many compatible functions that all operate on a common data structure.

`DIMPLE`supports the testing of various patient-level hypotheses and the exploration of heterogeneous tissue environments, by allowing the investigation of distance matrices in different parts of the tissue environment associated with varying levels of the intensity of a particular cell type. However, there are some limitations to the use of this software package. The smoothing bandwidth `bw` must be chosen a priori and can have a nontrivial influence on downstream results. We urge users to choose a biologically plausible “sphere of influence” when choosing the smoothing bandwidth. Furthermore, the individual cell-type frequencies may confound the association between patient outcomes and cell-cell interactions. Users may extract cell type counts to use in downstream modeling to adjust for cell specific overall intensities.

In the future, we plan to add support for incorporating information on other spatially-varying covariates, such as the levels of various cytokine or chemokine markers throughout the image space. We also aim to add support for incorporating various functional markers on different cells, such as Ki67 (a proliferation marker) or LAG3 (an immune checkpoint marker). These improvements will enhance the capabilities of `DIMPLE` and enable researchers to gain further insight into the relationships between different cell types in MI experiments, and is currently outside the scope of the current paper. Finally, we hope to add to our suite of open-source tools as we continue to develop statistical methods for the unique challenges presented by MI data.

## 5. Data and Code Availability

The `DIMPLE` R package is open source and freely available on Github (https://github.com/nateosher/DIMPLE). The Shiny app is available here: (https://bayesrx.shinyapps.io/DIMPLE_Shiny/).

The data used as illustration of our package was sourced from the `VectraPolarisData` package. This package can be installed from Bioconductor (https://bioconductor.org/packages/release/data/experiment/html/VectraPolarisData.html).

## 6. Acknowledgments

We thank Dr. Julia Wrobel for making available the data that is used in this manuscript. This work was supported by R37 CA214955-01A1 (MM,JE,VB,AR), P30 CA046592 (AR, MM, VB), R01 CA244845-01A1 (VB), T32 CA140044 (JE).

## 7. Author Contributions

M.M., N.O., and J.E. developed the presented software and drafted the manuscript. V.B. provided inputs in writing, reviewing and editing the manuscript. All authors contributed to conception of the project and approved the final version of the manuscript.

