## Supplement for "DIMPLE: AN R PACKAGE TO QUANTIFY, VISUALIZE, AND MODEL SPATIAL CELLULAR INTERACTIONS FROM MULTIPLEX IMAGING WITH DISTANCE MATRICES"

### 1. Loading and processing example data

In this section we will provide code which loads and pre-processes the example data in the manuscript. The data is publicly available lung cancer data from the `VectraPolarisData` package. This package can be installed from Bioconductor as follows (note that this code chunk, if run, will install the `BiocManager` package if it is not already installed):

```
if (!require("BiocManager", quietly = TRUE))
    install.packages("BiocManager")

BiocManager::install("VectraPolarisData")
library(VectraPolarisData)
lung_data = VectraPolarisData::HumanLungCancerV3()
lung_data
```

```
## dim: 8 1604786
## metadata(1): clinical_data
## assays(3): intensities nucleus_intensities membrane_intensities
## rownames(8): cd19_opal_650 cd3_opal_520 ... dapi autofluorescence
## rowData names(0):
## colnames: NULL
## colData names(124): cell_id tissue_category ... phenotype_cd4 sample_id
## reducedDimNames(0):
## mainExpName: NULL
## altExpNames(0):
## spatialCoords names(2) : cell_x_position cell_y_position
## imgData names(0):
```

Cell phenotype, is encoded in this data set by the named features `phenotype_cd14`, `phenotype_cd19`, `phenotype_cd4`, `phenotype_cd8`, `phenotype_ck`, `phenotype_other`

The phenotype variables are binary variables that indicate the type of each cell. We'll collect these variables into a single factor variable.

```
# Get the x coordinates for the cells
cell_x_coords = SpatialExperiment::spatialCoords(lung_data)[,1] %>% as.numeric()
# Get the y coordinates for the cells
cell_y_coords = SpatialExperiment::spatialCoords(lung_data)[,2] %>% as.numeric()
# Get the slide ids for the slides
slide_ids = lung_data$sample_id

# Map phenotypes to character vector - due to the size of
# the data, we take advantage of matrix multiplication methods
# to quickly map the six phenotype vectors to a single
# vector, or NA.
cell_marks = tibble(
```

```
  cd_14 = lung_data$phenotype_cd14[1:test_n] == "CD14+",
  cd_19 = lung_data$phenotype_cd19[1:test_n] == "CD19+",
  cd_4 = lung_data$phenotype_cd4[1:test_n] == "CD4+",
  cd_8 = lung_data$phenotype_cd8[1:test_n] == "CD8+",
  ck = lung_data$phenotype_ck[1:test_n] == "CK+",
  other = lung_data$phenotype_other[1:test_n] == "Other+"
) %>%
  as.matrix() %>%
  (\(m){
    m %*% 1:6
  }) %>%
  map_chr(\(x) c(NA, "CD14", "CD19", "CD4", "CD8", "CK", "Other")[x + 1])
```

Note that some cells don't have recorded phenotypes; we'll call those missing, and drop them for now.

```
  na_marks = which(is.na(cell_marks))
  cell_x_coords = cell_x_coords[-na_marks]
  cell_y_coords = cell_y_coords[-na_marks]
  slide_ids = slide_ids[-na_marks]
  cell_marks = cell_marks[-na_marks]
```

The patient level data for these slides can be extracted from the `lung_data` object as follows:

```
lung_metadata = metadata(lung_data)$clinical_data %>%
  tibble()
glimpse(lung_metadata)
```

```
## Rows: 153
## Columns: 13
## $ patient_id                <chr> "#01 0-889-121", "#02 1-037-393", "#...
## $ gender                    <chr> "F", "M", "M", "M", "M", "M", "F", "...
## $ mhcII_status              <chr> "low", "low", "high", "low", "high",...
## $ age_at_diagnosis          <dbl> 85, 66, 84, 79, 68, 57, 79, 73, 61, ...
## $ stage_at_diagnosis        <chr> "IA", "IA", "IIIA", "IA", "IA", "IA"...
## $ stage_numeric             <dbl> 1, 1, 3, 1, 1, 1, 2, 1, 2, 3, 2, 1, ...
## $ pack_years                <dbl> 60, 30, 50, 40, 20, NA, 40, 0, 35, 4...
## $ survival_days             <dbl> 3488, 1605, 176, 2042, 3747, 3443, 2...
## $ survival_status           <dbl> 0, 1, 1, 0, 0, 0, 1, 1, 1, 0, 0, 0, ...
## $ cause_of_death            <chr> NA, "Lung Ca", NA, NA, NA, NA, NA, "...
## $ adjuvant_therapy          <chr> "No", "No", "No", "No", "No", "No", ...
## $ time_to_recurrence_days   <dbl> NA, 1539, 176, NA, NA, NA, 285, 1740...
## $ recurrence_or_lung_ca_death <dbl> 0, 1, 1, 0, 0, 0, 1, 1, 1, 1, 0, 0, ...
```

In the raw data set, the patient identifiers are called `slide_id`; we have renamed them to `patient_id` for clarity. Something you might notice about the patient level data is that while there are 761 biopsies in the data set, there only seem to be 153 patients- thus, some patients have multiple slides of observation (in fact, virtually all patients have 5). In order to add metadata to a `MltplxExperiment` object, we have to include a `slide_id` variable in the metadata so that each row of the metadata can be linked to a slide in the object. When there is only one slide per patient, this can also function as a patient id, and no modification will be necessary; but in this case, because there are multiple slides per patient, we'll have to modify the metadata slightly before we attach it. To do so, we'll take advantage of the fact that the patient ID for each slide is the first 13 characters of the slide id:

```
slide_id_tibble = tibble(
  slide_id = unique(slide_ids),
  patient_id = substr(slide_id, 1, 13)
```

```
)

glimpse(slide_id_tibble)
```

```
## Rows: 761
## Columns: 2
## $ slide_id   <chr> "#01 0-889-121 P44_[40864,18015].im3", "#01 0-889-121...
## $ patient_id <chr> "#01 0-889-121", "#01 0-889-121", "#01 0-889-121", "#...
```

```
full_lung_metadata = left_join(
  slide_id_tibble,
  lung_metadata
)

glimpse(full_lung_metadata)
```

```
## Rows: 761
## Columns: 14
## $ slide_id                      <chr> "#01 0-889-121 P44_[40864,18015].im3...
## $ patient_id                    <chr> "#01 0-889-121", "#01 0-889-121", "#...
## $ gender                        <chr> "F", "F", "F", "F", "F", "M", "M", "...
## $ mhcII_status                  <chr> "low", "low", "low", "low", "low", "...
## $ age_at_diagnosis              <dbl> 85, 85, 85, 85, 85, 66, 66, 66, 66, ...
## $ stage_at_diagnosis            <chr> "IA", "IA", "IA", "IA", "IA", "IA", ...
## $ stage_numeric                 <dbl> 1, 1, 1, 1, 1, 1, 1, 1, 1, 1, 3, 3, ...
## $ pack_years                    <dbl> 60, 60, 60, 60, 60, 30, 30, 30, 30, ...
## $ survival_days                 <dbl> 3488, 3488, 3488, 3488, 3488, 1605, ...
## $ survival_status               <dbl> 0, 0, 0, 0, 0, 1, 1, 1, 1, 1, 1, 1, ...
## $ cause_of_death                <chr> NA, NA, NA, NA, NA, "Lung Ca", "Lung...
## $ adjuvant_therapy              <chr> "No", "No", "No", "No", "No", "No", ...
## $ time_to_recurrence_days       <dbl> NA, NA, NA, NA, NA, 1539, 1539, 1539...
## $ recurrence_or_lung_ca_death <dbl> 0, 0, 0, 0, 0, 1, 1, 1, 1, 1, 1, 1, ...
```

### 2. Simulating Data

The final category of functionality available in the DIMPLE package is the ability to easily simulate MI data in arbitrary patterns and intensities. First, we must create an "intensity grid." An "intensity grid" is a matrix that represents an underlying rectangular window. Each entry of the matrix represents a section of the underlying window of $k$ square units. It's most intuitive to take $k = 1$, but in principle $k$ can be any positive number. The actual number in the entry represents the intensity of cells of a certain type in that region; when $k = 1$, this is just the expected number of cells in that region. Here's an example intensity grid, and what it looks like when

```
grid_1 = GridRect(m = 100,
           n = 100,
           bot_left_corner_x = 10,
           bot_left_corner_y = 10,
           width = 50,
           height = 50,
           intensity = 0.1)

plot_simulation_heatmap(grid_1, t = "Example Intensity Grid")
```

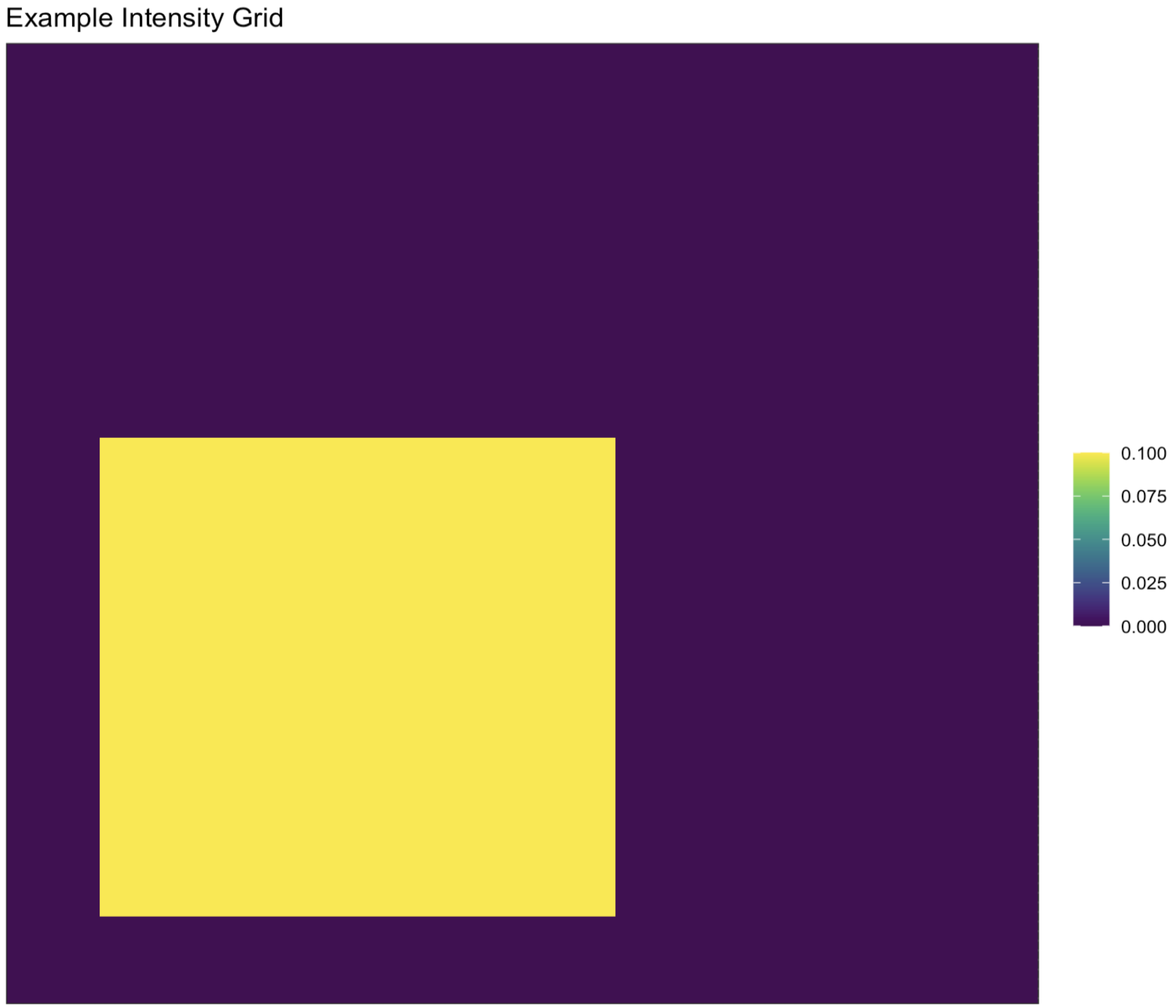


Note that `grid_1` is merely a `matrix` object. `GridRect` is one of two helper functions to generate such embeddings of geometric shapes within a larger matrix. The arguments `m` and `n` specify the width and height of the larger area in which the rectangle is embedded. `bot_left_corner_x` and `bot_left_corner_y` specify where, as you might expect, the x- and y- position of the bottom left hand corner of the rectangle. Note that the "origin" is considered to be the bottom left hand corner in order to be consistent with the coordinate system convention used by `spatstat`; thus, specifying the lower left-hand corner to be at (10, 10) indicates that the corner is located in the bottom left corner of the overall matrix, rather than the top-left. The `width` and `height` parameters naturally specify the width and height of the resulting rectangle; here we have set both to 50, resulting in a square intensity pattern. Finally, we specify that the intensity on this particular rectangle should be 0.1, i.e. we expect there to be about 1 point for every three by three grid of squares.

Once the intensity grid is generated, simulating a data set from it is trivial:

```
sim_1 = SimulateGrid(
  list(grid_1),
  square_side_length = 1
)

print(sim_1)
```

```
plot(sim_1)
```

```
## MltplxObject
## Slide id: PGWGNI
## Image with 271 cells across 1 cell types
## Cell types: Type 1
## No intensity generated (yet)
## No distance matrix generated (yet)
## 0 quantile distance arrays generated.
```

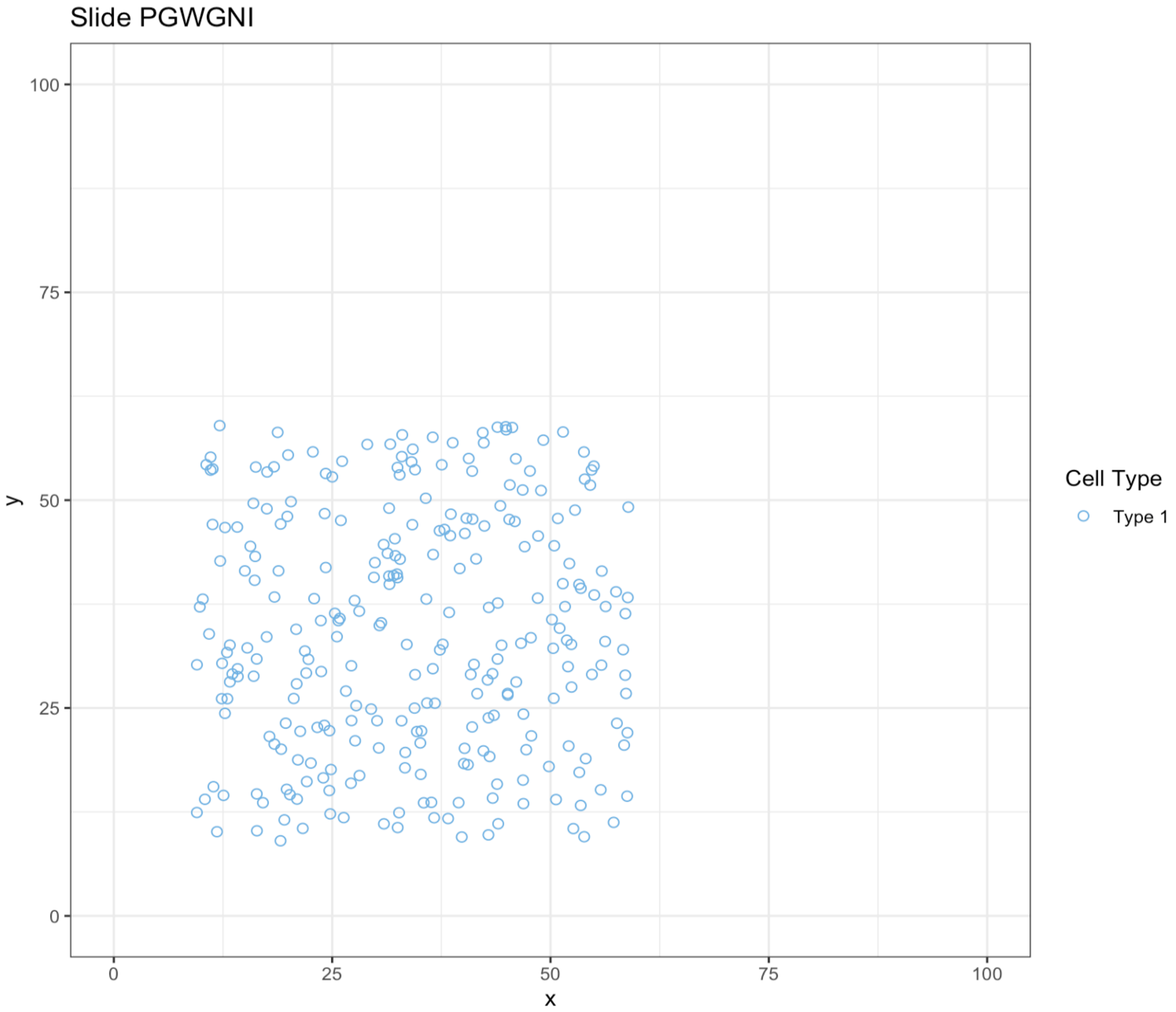


The `SimulateGrid` function takes a list of grids with common dimensions and simulates a point pattern with the specified intensity. It returns a `MltplxObject` with a randomly generated id. The `square_side_length` argument corresponds to the $k$ we discussed earlier; we specified it to be 1 for the sake of demonstration, but this is the default value, and thus can be omitted to achieve the same result.

The reason `SimulateGrid` takes a list of grids rather than a single grid is to simplify generating point processes of multiple types. For example, suppose we also generated an intensity grid for a point of a different type:

```
grid_2 = GridCircle(m = 100,
                    n = 100,
                    cx = 70,
                    cy = 70,
                    r = 25,
                    intensity = 0.1)

plot_simulation_heatmap(grid_2, t = "Example Intensity Grid 2")
```

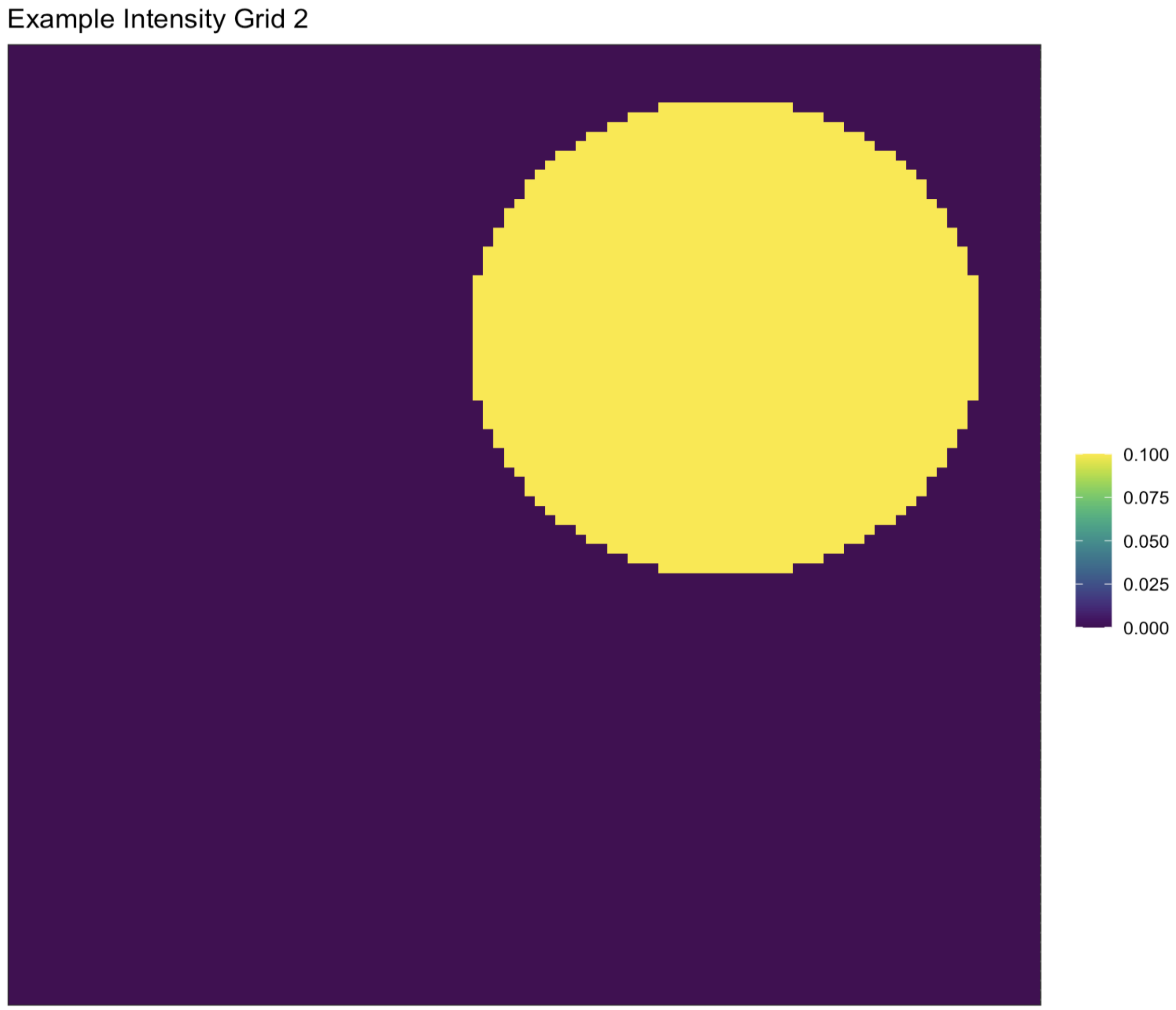


First, note that we have generated this intensity grid using `GridCircle`. The arguments are similar to `GridRect`, except that `cx` and `cy` specify respectively the x- and y-coordinates of the center of the circle, and `r` specifies its radius. Having generated these two intensity grids, it's trivial to simulate a point process with two types, which we'll label "Tumor" and "Immune" using the `marks` argument of `SimulateGrid`:

```
sim_2 = SimulateGrid(list(grid_1, grid_2),
                     marks = c("Tumor", "Immune"),
                     square_side_length = 1)

print(sim_2)
```

```r
plot(sim_2)
```

```
## MltplxObject
## Slide id: OGBQZP
## Image with 460 cells across 2 cell types
## Cell types: Immune, Tumor
## No intensity generated (yet)
## No distance matrix generated (yet)
## 0 quantile distance arrays generated.
```

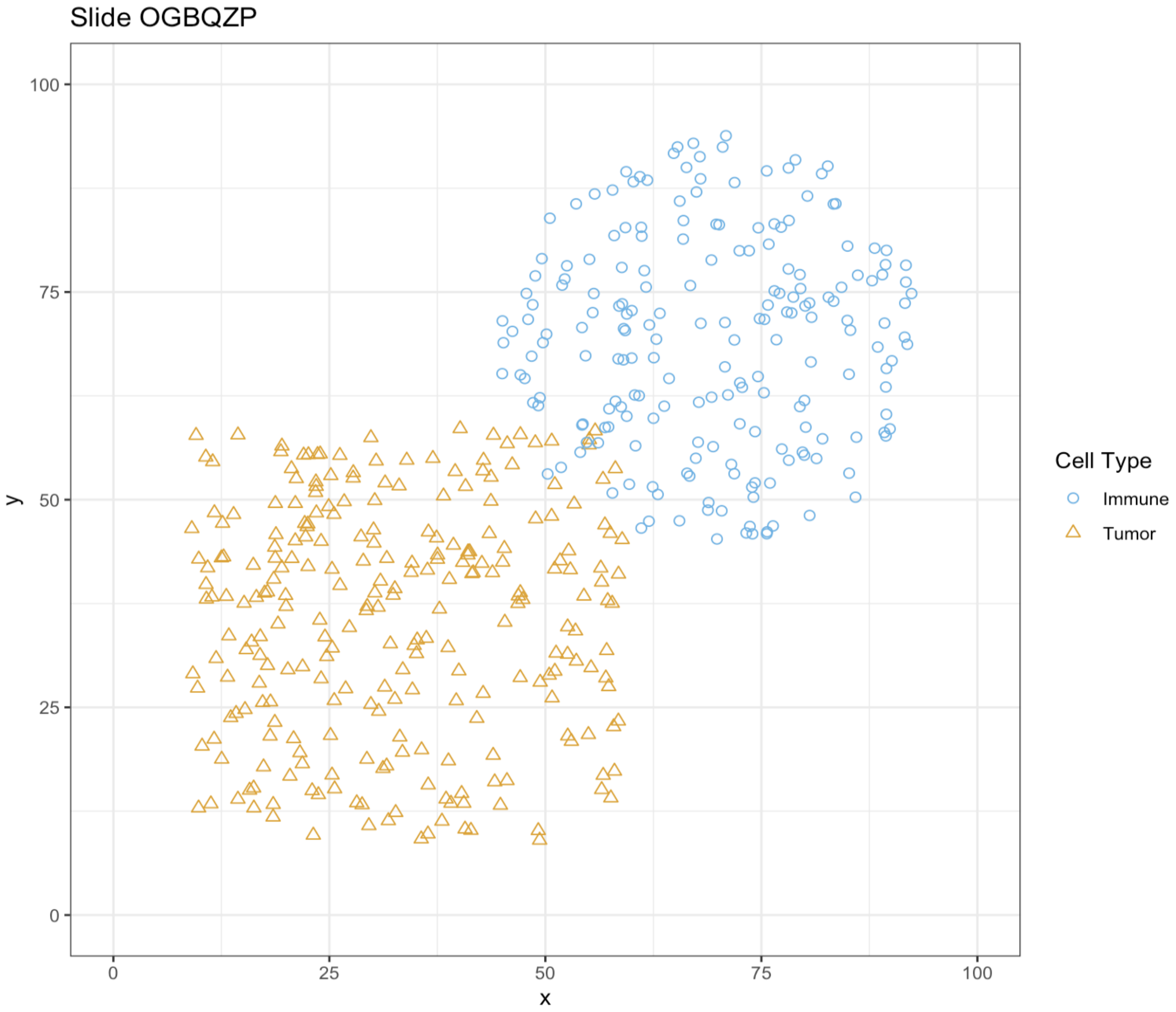


We conclude by noting that individual `MltplxObjects` generated from simulations can be gathered into a list and converted to a `MltplxExperiment` using the `as_MltplxExperiment` generic provided by `DIMPLE`:

```r
sim_exp = list(sim_1, sim_2) %>%
  as_MltplxExperiment()

print(sim_exp)
```

```
## MltplxExperiment with 2 slides
## No intensities generated
## No distance matrices generated
## No attached metadata
```
